# VLCFA-mediated inter-cell layer communication controls cellular pluripotency in Arabidopsis callus

**DOI:** 10.64898/2026.09.01.748728

**Authors:** Yuki Doll, Takashi Nobusawa, Mikiko Kojima, Kenji Nagata, Kanae Matsuda-Ito, Ari Pekka Mähönen, Taito Matsuda, Mitsutomo Abe, Hitoshi Sakakibara, Momoko Ikeuchi

## Abstract

Plants have remarkable capacity to reconstruct entire organ systems from tissue explants. In Arabidopsis two-step tissue culture system, pluripotency regulators are specifically expressed in the middle-cell layer of the stratified callus tissue. However, regulatory mechanisms underlying the radial patterning of callus remained unclear. Here, we found that very-long-chain fatty acids (VLCFAs) synthesized in the epidermis-like outermost layer are essential for pluripotency acquisition and successful shoot regeneration. Our genetic and transcriptomic analyses revealed that the regulatory roles of VLCFAs on pluripotency acquisition involve inter-cell layer signaling in callus tissue, while they are at least partly independent of ATML1/PDF2 functions and cuticular wax synthesis in the outermost layer. VLCFAs spatially restrict procambium cell identity by non-cell-autonomously suppressing cytokinin signaling, thereby allowing for establishment of the middle-cell layer. We propose that the inhibitory relationships between layer-specific regulators underlie the intricate balance of cellular fate determination in pluripotent callus.

## INTRODUCTION

Plants display astonishing regenerative responses to newly produce entire organ systems from tissue explants. During *de novo* organogenesis, shoot or root stem cell niche is re-established, often via production of pluripotent cell mass called callus (*1*, *2*). Molecular and developmental basis of *de novo* organogenesis has been well characterized using *Arabidopsis thaliana* (Arabidopsis) tissue culture. In the Arabidopsis two-step culture system, root or hypocotyl explants pre-cultured on auxin-rich callus inducing media (CIM) develop pluripotent callus from xylem-pole pericycle (XPP) cells (*3*, *4*). The pluripotent callus is then transferred to the shoot inducing media (SIM) or root inducing media (RIM) to induce *de novo* formation of shoots or roots, respectively (*1*, *2*).

Pluripotent callus formed during pre-culture on CIM, which we term here “CIM callus”, has a well-organized layered structure that resembles incipient root primordia. Root quiescent center (QC) regulators including SCARECROW (SCR) and WUSCHEL-RELATED HOMEOBOX 5 (WOX5) are specifically expressed in the middle-cell layer of the CIM callus and confers cellular pluripotency (*5*, *6*). Single-cell RNA-seq and imaging analyses revealed characteristic gene expression profiles in other cell layers as well. Epidermis markers including *ARABIDOPSIS THALIANA MERISTEM L1 LAYER* (*ATML1*) are specifically expressed in the outermost layer of the CIM callus (*5*). Inner cell layers of the callus are characterized by a set of marker genes such as *SHORTROOT* (*SHR*) and *TARGET OF MONOPTEROS5* (*TMO5*) (*5*, *6*). Upon transfer to SIM for shoot induction, callus develops into a more heterogenous cell population, which we term here “SIM callus.” Subpopulation of the middle-cell-layer cells in CIM callus are specified into shoot apical meristems (*5*). Similar to CIM callus, SIM callus also harbors the outermost layer with *ATML1* expression and inner layers with procambial marker expression (*7*). Although accumulating evidence provides cell layer-specific gene expression profiles, it remains unknown how this radial spatial pattern is established *de novo*, and whether regulators in each specific layer have independent or inter-connected regulatory relationships.

Inter-cell layer communications via mechanical and biochemical signals constitute key regulatory aspects of plant development. During shoot development, the epidermal tissue physically constrains growth of the inner tissue layers and coordinates organ growth (*8*, *9*). Shoot epidermis also produces various kinds of signaling molecules such as peptides and microRNAs that non-cell autonomously regulate gene regulation in inner layers (*10–12*). Fatty acids with more than 18 carbons, called very-long-chain fatty acids (VLCFAs), are synthesized in the epidermis and serve as precursors for cuticular waxes and various lipids (*13*). They are synthesized in the endoplasmic reticulum from long-chain fatty acids (LCFAs) via stepwise elongation of the carbon chain, a process catalyzed by the fatty acid elongase (FAE) enzyme complex. Apart from their structural functions, VLCFAs or their derivatives also mediate inter-cell layer communications in various developmental processes (*14–20*). For instance, VLCFA synthesis in the shoot epidermis is required for the proper control of vascular cell proliferation (*14*). VLCFA is also crucial for the plasma-membrane localization of auxin transporters PIN-FORMEDs (PINs) in roots or in embryos (*15*, *20*). Furthermore, VLCFAs constitute a positive feedback loop with master regulators of epidermis identity specification, ATML1 and PROTODERMAL FACTOR 2 (PDF2), via physical interaction and stabilization of the protein complexes (*21*, *22*). Several lines of evidence suggest the involvement of VLCFAs in callus formation and shoot regeneration (*16*, *17*, *23*, *24*); however, the underlying mechanisms remain elusive.

In this study, we find that VLCFA synthesis in the outermost layer of Arabidopsis callus spatially restricts procambium domain to inner layers, thereby facilitates the establishment of the middle-cell layer that endows pluripotency to callus cells. Our study highlights the importance of inter-layer communication in pluripotency acquisition and organ regeneration.

## RESULTS

### VLCFA synthesis is required for shoot regeneration from hypocotyl and root explants

In our attempt to characterize cell layer-specific factors in shoot regeneration, we tested potential roles of VLCFAs, whose synthetic genes are specifically expressed in epidermis-like cell clusters in single cell RNA-seq data from both CIM and SIM calli (*5*, *7*) (fig. S1). We treated Arabidopsis explants with cafenstrole (CF), a chemical that blocks the first step of VLCFA chain elongation catalyzed by FAE component ketoacyl-CoA synthases (KCSs) (Fig. 1A) (*25*). Strikingly, CF treatment dose-dependently inhibited shoot regeneration from hypocotyl explants (Fig. 1B−C). Gas chromatographic analyses verified that VLCFA content was significantly reduced upon 0.3 μM CF treatment (fig. S2A). Similar to CF, another KCS inhibitor metazachlor (Meta) (*26*) had inhibitory effects on shoot regeneration as well (fig. S2B, C). Our genetic analyses further revealed that FAE component mutants *cer10-2* (*27*), *kcs1-5* (*17*), and *pas2-1* (*28*) are defective in shoot regeneration (Fig. 1A, D, E and fig. S2D, E). Not only hypocotyls but also root explants exhibited severe defects in shoot regeneration in VLCFA deficient conditions, including CF-treated wild-type explants, *kcs1-5*, and an FAE scaffold mutant *pas1-4* (*17*), corroborating the general requirement of VCLFA synthesis in shoot regeneration (fig. S2F−K). We also confirmed that shoot regeneration defect of *kcs1-5* was rescued by introducing *pKCS1:KCS1-GFP* (fig. S2L−M), before using this mutant for subsequent analyses. These results together demonstrate that VLCFA synthesis is necessary for shoot regeneration in the two-step culture system.

**Figure 1.**
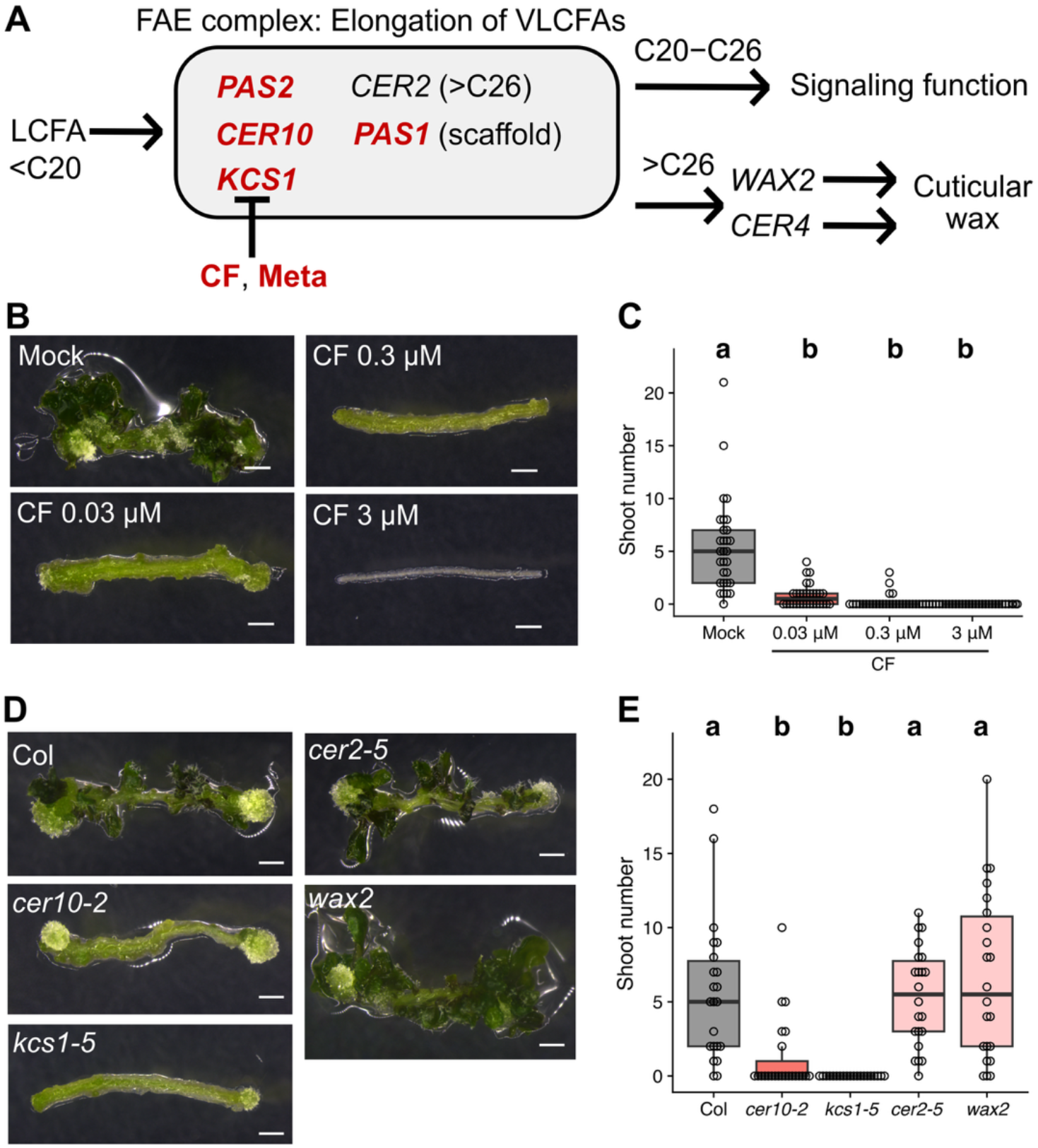
VLCFA synthesis is necessary for shoot regeneration in the two-step culture system. (A) Schematic diagram of the VLCFA synthesis pathway, with the information of genes and inhibitors analyzed in this study. Genes or inhibitors are highlighted in red if respective mutations or inhibitor treatment caused severe regeneration defects as shown in Fig.1 and Fig. S2. (B–C) Shoot regeneration phenotype of wild-type hypocotyl explants under continuous cafenstrole (CF) treatment. Different letters indicate statistically significant differences (*p* < 0.05 in Tukey-Kramer test, *n* = 30). (D–E) Shoot regeneration phenotypes of VLCFA-related mutant hypocotyl explants. Different letters indicate statistically significant differences (*p* < 0.05 in Tukey-Kramer test, *n* > 19). Bars: 1 mm.

VLCFAs with longer carbon chains (> C26) serve as precursors for cuticular wax synthesis, whereas those with shorter carbon chains (< C28) are incorporated into intracellular lipids and putatively act as a signaling molecule (*13*, *14*). Using genetic approaches, we sought to identify which type of VLCFAs has critical roles in shoot regeneration. We analyzed a mutant of *ECERIFERUM 2* (*cer2-5*) (*29*, *30*), an FAE component that is responsible for elongating VLCFAs to exceptional lengths (> C26), and mutants of genes involved in the processes synthesizing cuticular waxes from exceptionally long VLCFAs (*wax2*, *cer4-1*) (*31*, *32*). We found that all these mutants regenerated shoots with no discernable defect (Fig. 1A, D, E and fig. S2N, O), suggesting that longer VLCFAs and cuticular waxes synthesized from such VLCFAs are dispensable for shoot regeneration. Therefore, we conclude that shorter VLCFAs (C20−C26) are specifically required for successful shoot regeneration.

### VLCFAs are synthesized in the newly established epidermal tissue of callus

Next, we asked where and when VLCFAs are produced during the two-step tissue culture of hypocotyl explants. To trace the spatiotemporal dynamics of VLCFA synthesis along with the establishment of epidermal tissue, we analyzed the expression pattern of an FAE component enzyme PAS2 (*33*) together with the epidermis marker ATML1. Before CIM incubation, the expression of PAS2 and *ATML1* are not detected in the dark-grown hypocotyl explant, except occasional *ATML1* expression in the hypocotyl epidermis (Fig. 2A). It is noteworthy that PAS2 is not expressed in any inner tissues in hypocotyls, while we confirmed its endodermal expression in roots (fig. S3A), as previously reported for this enzyme and other FAE components (*17*, *22*). After 1 or 2 days of pre-incubation on CIM (CIM1−2D), the expression of both genes initiated in the emerging callus cells, specifically in the outer cells produced by periclinal divisions of pericycle cells (Fig. 2A and fig. S3B). This expression onset after periclinal division is similar to that observed during lateral root initiation (*22*). As callus matures, callus cells continue to proliferate periclinally to produce multiple cell layers. At CIM4D, the expression of PAS2 and *ATML1* remained specific to the outermost layer of the callus (Fig. 2A). Upon transfer to SIM, the expression of PAS2 and *ATML1* gradually became specified to a subset of cells in the outermost layer (fig. S3B). The expression of both genes was restricted to callus cell populations with a smooth surface composed of relatively small polygonal cells, while it was absent from rough callus composed of globular expanded cells on SIM7D (Fig. 2A). This observation is consistent with our previous single-cell RNA-seq analysis and imaging analyses of the ATML1 reporter (*7*). Taken together, the epidermis-like tissue with ATML1 expression is newly established during shoot regeneration in a stepwise manner, where it first initiates as the outermost cell layer of CIM callus and then becomes restricted to the surface of meristematic cell populations in SIM callus. Our results indicate that VLCFAs are synthesized in this outermost cell layer throughout callus development and shoot regeneration (Fig. 2B).

**Figure 2.**
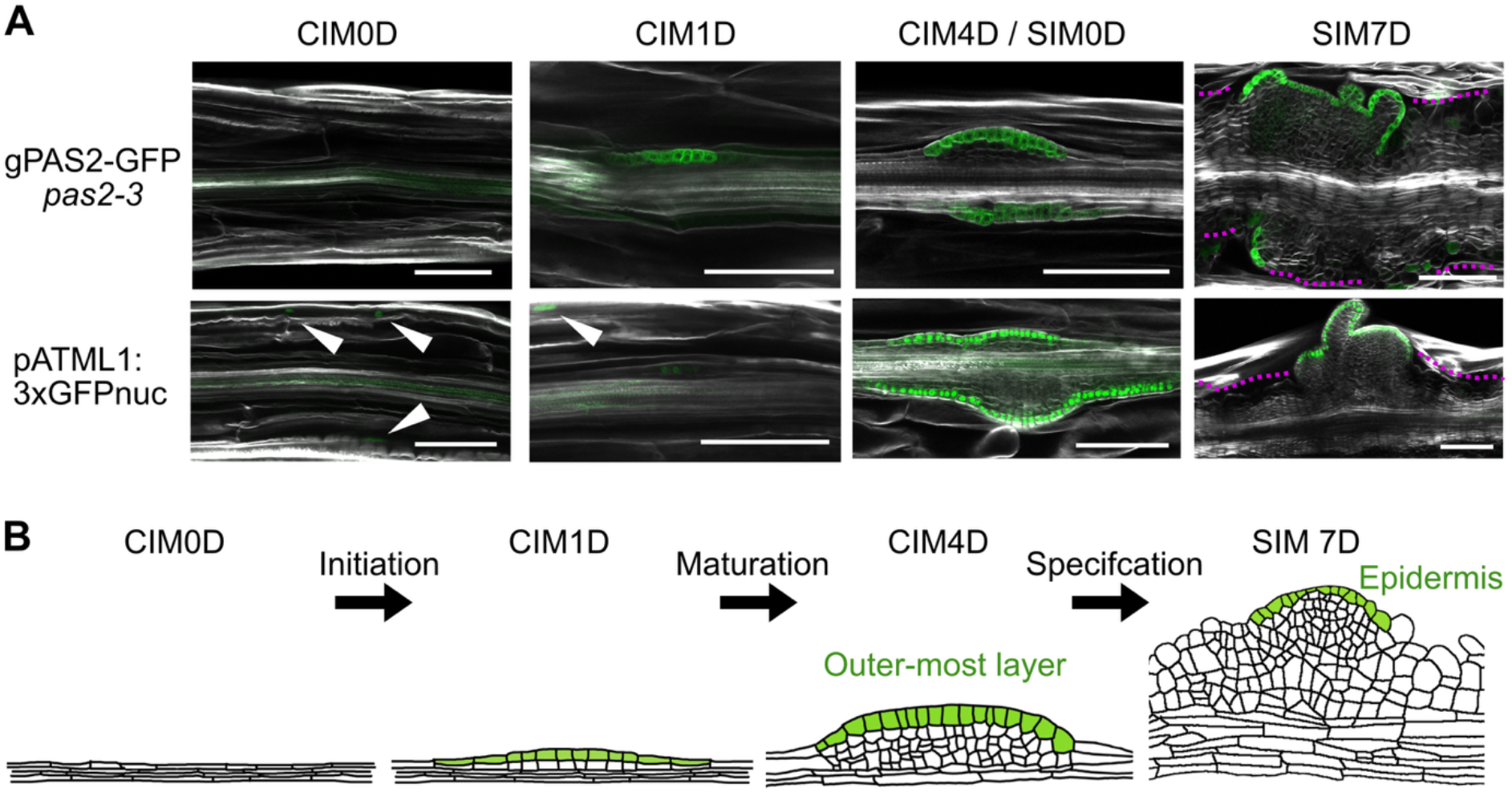
VLCFAs are synthesized in the re-established epidermal tissue during two-step culture. (A) Localization of fluorescence reporters during callus formation and shoot regeneration in the two-step tissue culture. Arrowheads indicate expression of *ATML1* in the original hypocotyl epidermis. Dotted magenta lines indicate callus regions without epidermal marker expression. Representative images from observation of at least 5 explants are shown. Bars: 100 μm. (B) Schematic image of the process of epidermal tissue re-establishment.

### VLCFA synthesis positively regulates callus maturation and pluripotency acquisition

To identify the specific incubation stage when VLCFAs have critical functions, we applied CF in a stage-specific manner during CIM or SIM incubation. Whereas CF treatment during SIM incubation mildly inhibited shoot regeneration, CF treatment during pre-incubation on CIM alone almost completely blocked shoot regeneration (Fig. 3A−B). This result indicates that VLCFA synthesis is important in both CIM and SIM stages, yet it has more profound effects during the CIM pre-culture stage. The expression of *WUSCHEL* (*WUS*), one of the earliest hallmark genes of shoot progenitor establishment (*34*), failed to be established in *kcs1-5* or CF-treated calli upon transfer to SIM (Fig. 3C), which is consistent with the idea that their defects originate from the earlier pre-incubation step. To further uncover the molecular basis of VLCFA effect, we analyzed the expression of SCR and WOX5, middle-cell layer-specific QC regulators that endow pluripotency to CIM callus (*5*, *6*). In the VLCFA-deficient callus, the expression of *WOX5* and SCR was weaker and spatially diffused to outer cell layers (Fig. 3D). Furthermore, VLCFA-deficient callus regenerated less adventitious roots after incubation on RIM (fig. S4A−D), together supporting the idea that VLCFAs are required for pluripotency acquisition of CIM callus.

**Figure 3.**
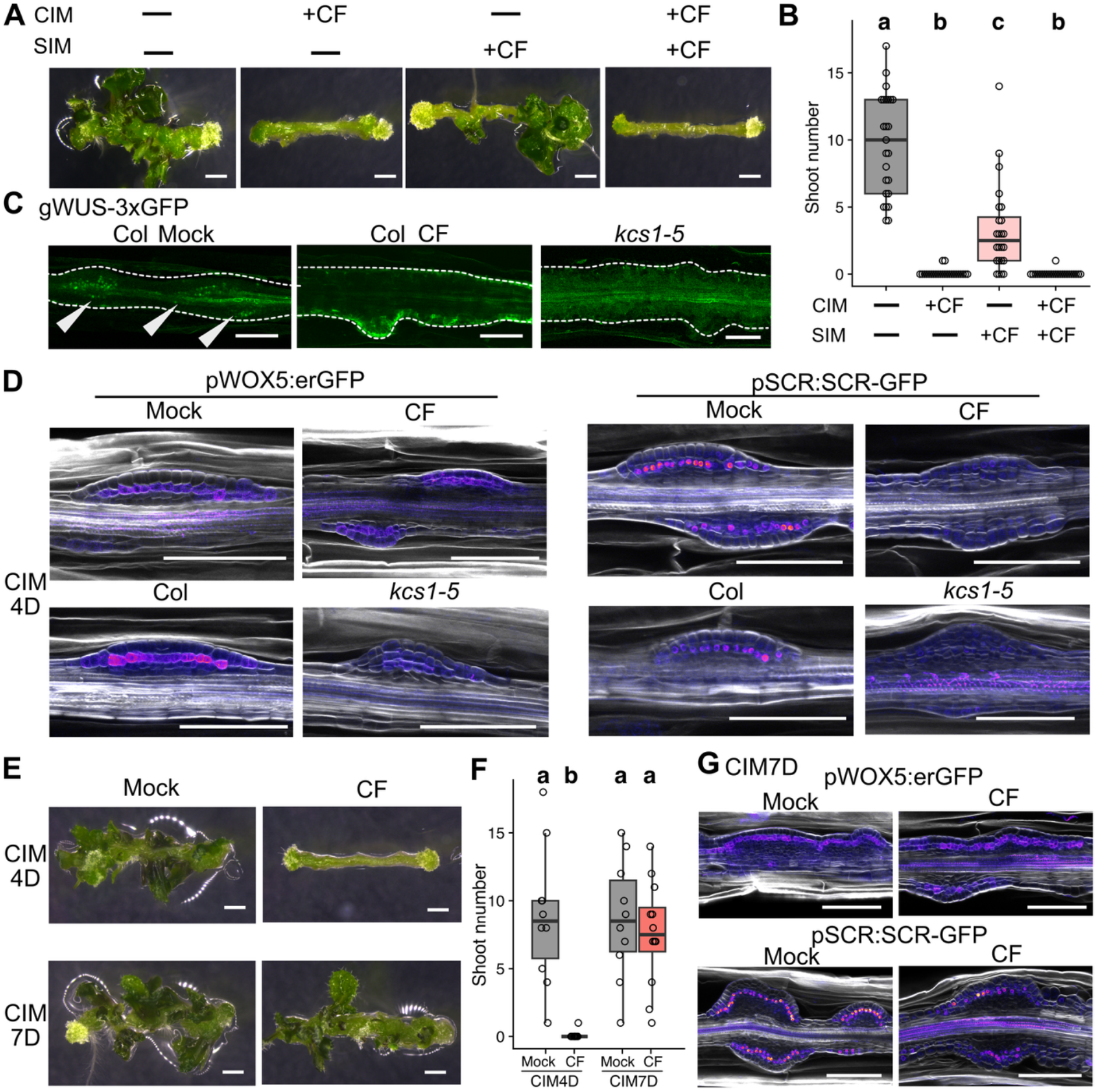
VLCFAs are needed for pluripotency acquisition of CIM callus. (A–B) Effect of stage-specific CF treatment on shoot regeneration from hypocotyl explants. B: Different letters indicate statistically significant differences (*p* < 0.05 in Tukey-Kramer test, *n* > 23). (C) Expression of WUS reporter in VLCFA-deficient callus during SIM culture at SIM4D. Arrowheads indicate groups of WUS expressing cells. (D) Expression of pluripotency markers *WOX5* and SCR on VLCFA-deficient callus at CIM4D. (E–F) Effect of prolonged CIM culture on shoot regeneration of CF-treated callus. Different letters in (F) indicate statistically significant differences (*p* < 0.05 in Tukey-Kramer test, *n* > 19). (G) Expression of pluripotency markers on VLCFA-deficient callus at CIM7D. Representative images in (C, D, G) are from observation of at least 5 explants Bars: 1 mm (A, E), 100 μm (C, D, G).

Notably, the observed weak and diffused expression of QC-related genes in the VLCFA-deficient callus resembled normal CIM callus at an earlier stage around CIM2D (*6*). This led us to speculate that callus maturation might be delayed in VLCFA-deficient callus. We therefore extended the pre-incubation on CIM and found that 7-day culture on CIM alleviated the shoot regeneration defect caused by CF treatment (Fig. 3E, F). Consistently, the expression of *WOX5* and SCR was localized to the middle-cell layer under CF treatment on CIM7D (Fig. 3G). These data suggest that VLCFAs non-cell autonomously regulate the establishment of QC-regulator expression in the middle-cell layer via promotion of callus maturation. Prolonged CIM culture, however, did not rescue shoot regeneration defects in VLCFA-deficient mutants (fig. S4E, F), probably because VLCFA deficiency during the SIM stage impairs regeneration in these mutants (Fig. 3A, B).

### VLCFA synthesis is necessary for establishing layer-specific gene regulatory networks

As our imaging analyses suggested profound and non-cell autonomous effects of VLCFA deficiency on callus gene expression, we sought to comprehensively explore downstream events of VLCFAs. We performed two sets of comparative transcriptome analyses on CIM4D: wild type vs. *kcs1-5* and CIM with vs. without CF. Our principal component analysis clearly separated control and VLCFA-deficient conditions from both sets of experiments (fig. S5A). Differentially expressed genes (DEGs) for the two comparisons substantially overlapped, indicating that our RNA-seq experiment successfully captured the VLCFA-dependent transcriptomic signature (fig. S5B). Among 682 common DEGs that were upregulated in the VLCFA-deficient conditions, Gene Ontology (GO) categories such as stress response and synthesis of phenylpropanoids including lignin were significantly enriched (Fig. 4A). Among 731 common DEGs that were downregulated in VLCFA-deficient conditions, genes involved in meristem development and epidermal development were highly enriched, which suggests important developmental roles of VLCFAs (Fig. 4A). Further examination of meristematic genes included in the DEGs revealed that many middle-cell layer-specific QC-related genes defined in the previous studies (*5*, *6*), including *WOX5* and *SCR*, were drastically downregulated in the VLCFA-deficient conditions (Fig. 4B). In addition, we detected clear downregulation of outermost layer-specific epidermal genes (*5*, *6*), including *ATML1* and *PDF2* (Fig. 4C). Consistently, we detected a clear decrease in translational ATML1 reporter accumulation in CF-treated callus (Fig. 4D), which is likely attributable to both lower transcription rate and protein destabilization by VLCFA deprivation (*22*). Genes transcribed in the inner layers of the callus (*5*, *6*), including *SHR* and *TMO5*, were also downregulated (Fig. 4E). We confirmed that the accumulation of fluorescent SHR protein in the inner- and middle-cell layers of callus clearly decreased under VLCFA deficiency (Fig. 4F). These findings corroborate the idea that the outermost layer-specific VLCFAs regulate the establishment of gene regulatory networks that are specifically active in three respective layers.

**Figure 4.**
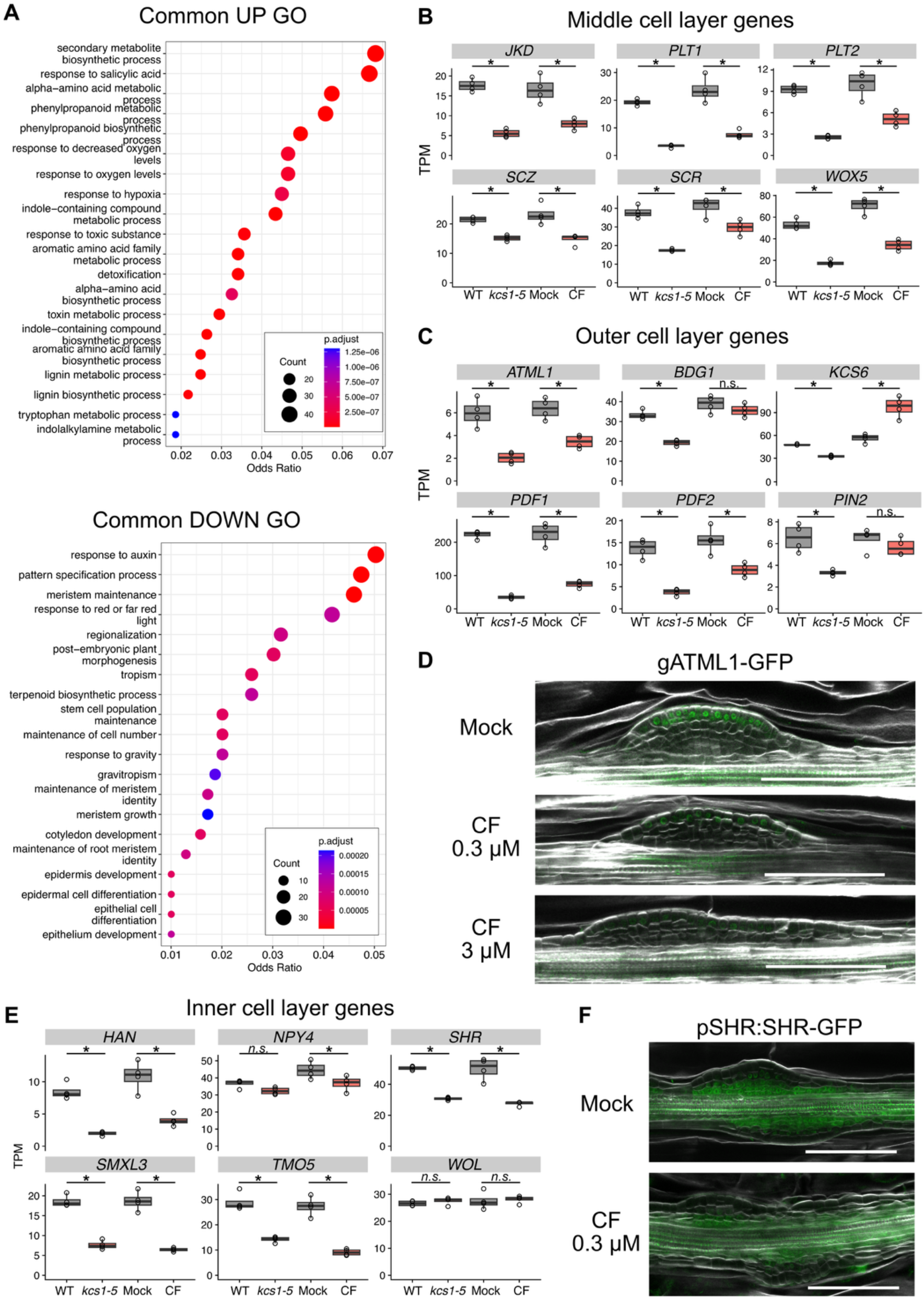
VLCFAs are needed for establishing layer-specific gene regulatory networks. (A) Results of GO enrichment analyses for common DEGs upregulated or downregulated in VLCFA-deficient conditions. (B–C) The expression of the middle cell layer-specific genes (B) and the outermost cell layer-specific genes (C) in the RNA-seq dataset. (D) Expression of translational ATML1 reporter gATML1-GFP in CIM callus under treatment with different concentrations of CF. (E) The expression of the inner cell layer-specific genes in the RNA-seq dataset. (F) Expression of SHR in CIM callus treated with CF. (D, F) Representative images from observation of at least 5 explants are shown. Bars: 100 μm. *: FDR < 0.05 in edgeR.

### ATML1/PDF2 are not likely the downstream of VLCFA function in CIM callus

Based on our transcriptomic analysis, we first hypothesized that the pluripotency defect in VLCFA-deficient conditions may be attributable to the down-regulation of *ATML1* and *PDF2*, which have been implicated in callus pluripotency by a recent study (*23*). Our phenotypic analyses revealed that *atml1-1 pdf2-1*, whose seedlings lack functional shoot epidermis (*35*), failed to regenerate any visible shoots either from hypocotyl or root explants (Fig. 5A and fig. S6), demonstrating essential roles of ATML1/PDF2 in shoot regeneration. In contrast to VLCFA-deficient conditions, however, the *atml1-1 pdf2-1* mutant showed no defect in layered tissue structure or expression of the pluripotency marker gene *WOX5* on CIM4D (Fig. 5B), indicating that ATML1/PDF2 may not be involved in pluripotency acquisition of CIM callus. In addition, WUS-expressing cells were properly generated in the mutant callus on SIM (Fig. 5C), suggesting that developmental processes leading to the establishment of early shoot progenitors are not affected in the *atml1-1 pdf2-1* mutant. Indeed, *atml1-1 pdf2-1* occasionally produced malformed SAMs with cylindrical leaf-like structures (Fig. 5A, D), which resembled the malformed leaves produced in *atml1-1 pdf2-1* seedlings (*9*, *35*). These observations indicate that *atml1-1 pdf2-1* mutant is defective in shoot organogenesis, but not in pluripotency acquisition of callus.

**Figure 5.**
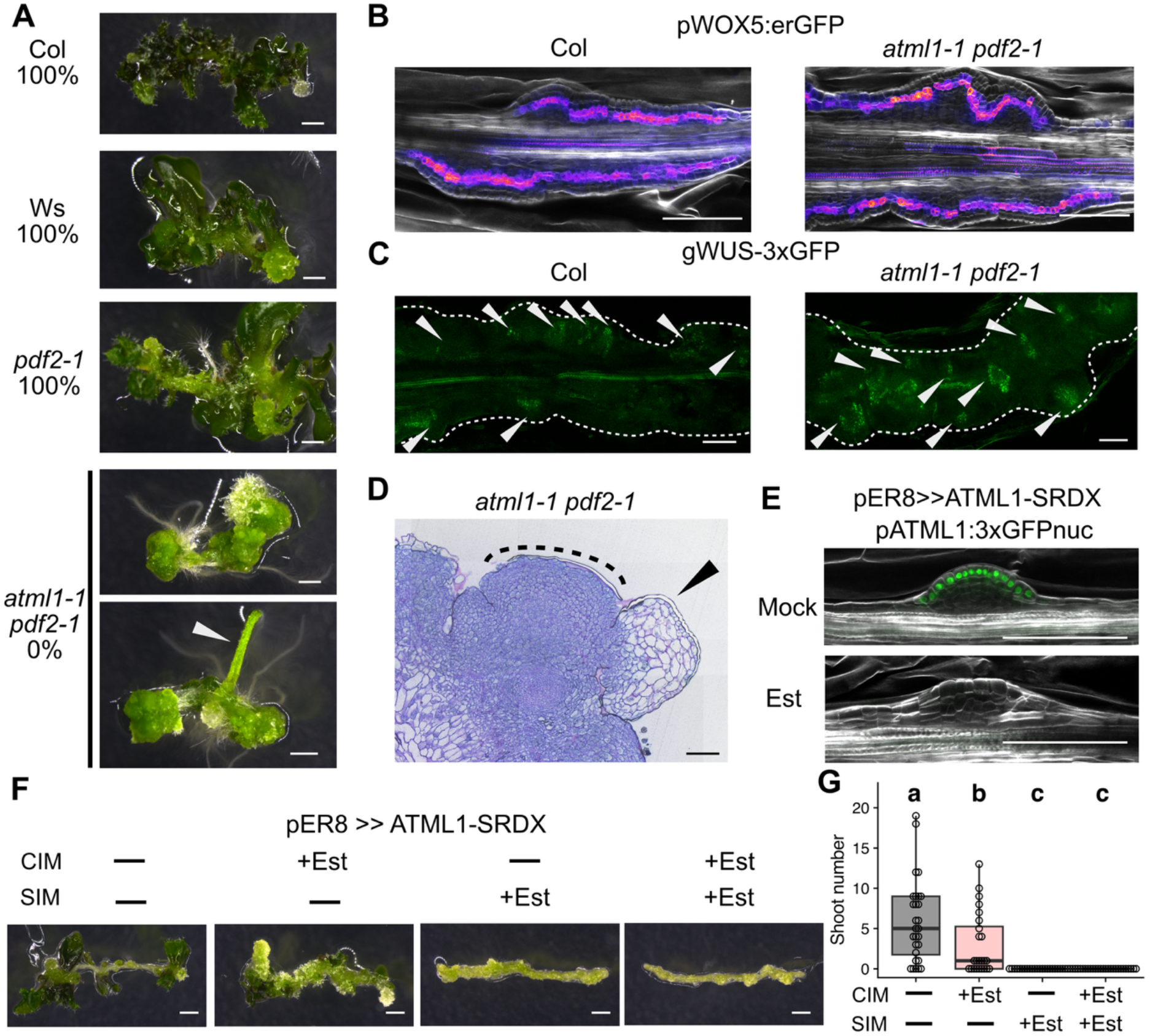
ATML1/PDF2 are likely dispensable for pluripotency acquisition and rather regulates shoot morphogenesis on SIM. (A) Shoot regeneration phenotype of *atml1-1 pdf2-1* double mutant hypocotyl explants. The arrowhead indicates a cylindrical leaf-like structure. The frequency of shoot regeneration is shown (*n* = 18 (Col), 16 (Ws), 26 (*pdf2-1*), 8 (*atml1-1 pdf2-1*)) (B) Expression of WOX5 reporter in *atml1-1 pdf2-1* at CIM4D. (C) Expression of WUS reporter in *atml1-1 pdf2-1* at SIM6D. Arrowheads indicate groups of WUS expressing cells. (D) Malformed SAM-like structure in *atml1-1 pdf2-1* (dotted line) with a leaf-like structure (arrowhead) at SIM10D. (E) Expression of *ATML1* reporter under ATML1-SRDX induction by 5 μM estradiol at CIM4D. (F–G) Effect of stage-specific induction of ATML1-SRDX on shoot regeneration. Representative images in (B, C, F) are from observation of at least 5 explants are shown. Different letters in (G) indicate statistically significant differences (*p* < 0.05 in Tukey-Kramer test, *n* > 23). Bars: 1 mm (A, G), 100 μm (B–F).

We should note, however, that *atml1-1* is a weak allele and thus phenotypic analyses of *atml1-1 pdf2-1* are not sufficient for drawing a conclusion about functional requirement of ATML1/PDF2 (*35*, *36*). As double null *atml1 pdf2* mutations result in embryonic arrest (*36*), we exploited an alternative strategy to chemically induce a chimera repressor ATML1-SRDX, which has strong dominant-negative effects on seedling growth (*37*). Induced expression of ATML1-SRDX abolished the *ATML1* promoter activity in CIM callus (Fig. 5E), as expected from the self-activating positive feedback regulation of ATML1 (*35*). However, our stage-specific induction experiment revealed that ATML1-SRDX induction during CIM culture has only minor effects on subsequent shoot regeneration (Fig. 5F, G), suggesting that ATML1 is likely dispensable for pluripotency acquisition, unlike VLCFAs. In contrast, ATML1-SRDX induction during the SIM stage severely inhibited shoot regeneration (Fig. 5F, G). These results together demonstrate that ATML1/PDF2 play important roles rather in the morphogenetic process during SIM culture to construct functional SAMs. We therefore conclude that regulatory roles of VLCFAs in pluripotency acquisition is not mediated by ATML1/PDF2.

### VLCFAs positively regulate pluripotency acquisition by suppressing cytokinin signaling in the callus inner layer

We then pursued the possibility that VLCFAs regulate pluripotency acquisition by controlling phytohormone dynamics of the callus cels, as our results so far suggested that VLCFAs non-cell-autonomously regulate middle- and inner-layer-specific gene expression (Fig. 4). Previous studies suggested that VLCFAs regulate auxin signaling by mediating localization of PIN auxin efflux carriers in multiple developmental contexts (*15*, *20*). However, we did not find any substantial differences in PIN1 localization in CIM callus with or without CF treatment, except that PIN1 accumulation level was slightly lower in the CF-treated callus (fig. S7A). Also, the expression of the auxin response reporter DR5rev:GFP was not affected by CF treatment (fig. S7B), together implying that auxin signaling is not likely the primary downstream of VLCFAs in pluripotency acquisition.

A previous study showed that VLCFAs derived from shoot epidermis suppress cytokinin biosynthesis in inner tissues (*14*). We therefore investigated the possibility that cytokinin may be involved in the VLCFA-mediated pluripotency acquisition. Under control conditions, the expression of a cytokinin response reporter TCSn:GFP (*38*) was restricted to the vascular region of the explant and was excluded from callus region (Fig. 6A). In contrast, under VLCFA-deficient conditions, the signal expanded into the inner layers of the callus and was occasionally detected even in the outermost layer (Fig. 6A). The RNA-seq data supported this observation, as several cytokinin-responsive type-A *ARABIDOPSIS RESPONSE REGULATOR* (*ARR*) genes (*39*) were upregulated in VLCFA-deficient conditions (fig. S8A). Collectively, these results suggest that VLCFA deficiency causes hyperactivation of cytokinin responses in CIM callus.

**Figure 6.**
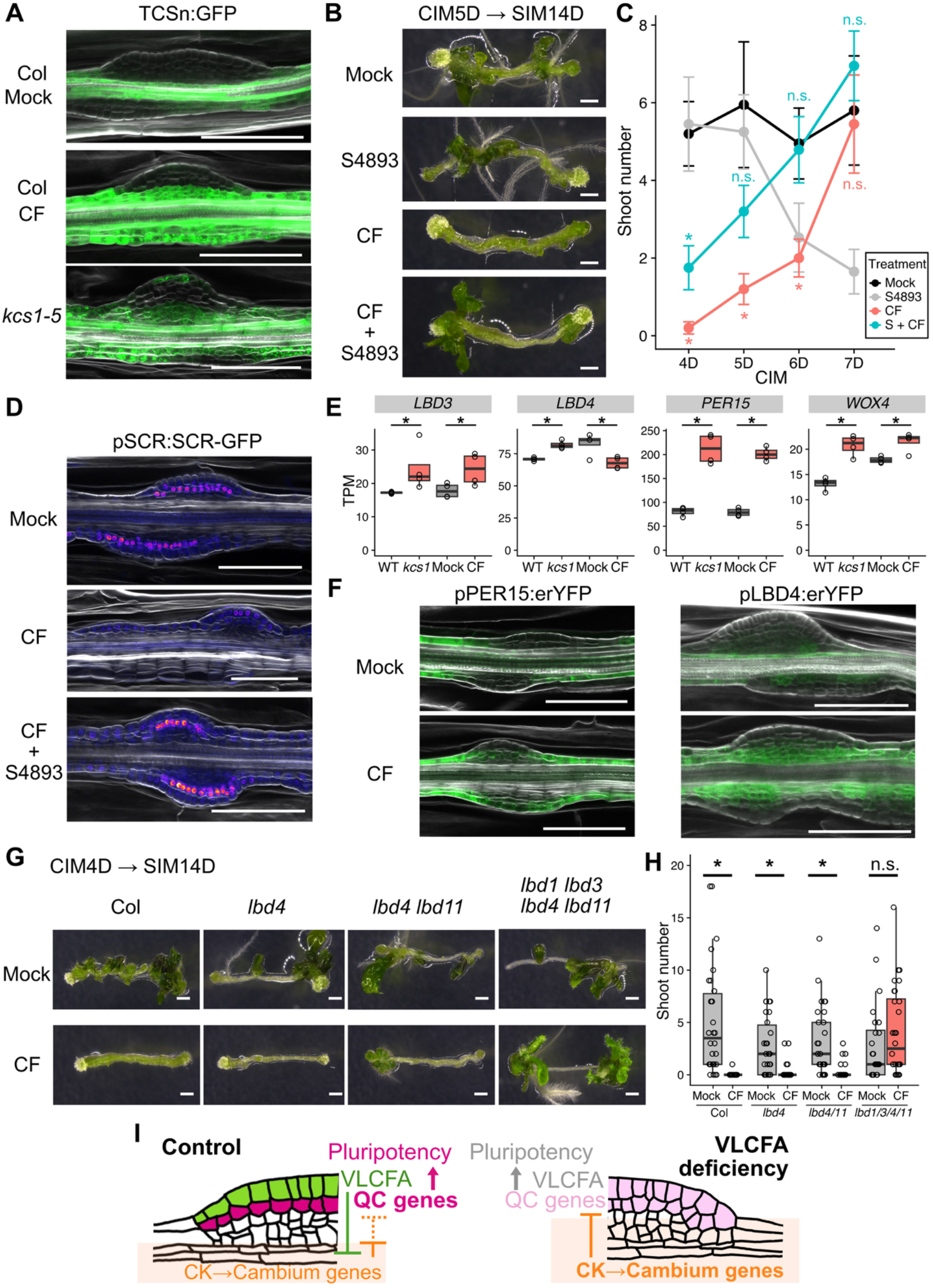
VLCFAs facilitate pluripotency acquisition by suppressing cytokinin signaling and procambium-like gene regulatory networks in the callus inner layer. (A) Expression of the cytokinin response reporter TCSn:GFP in VLCFA-deficient CIM callus. (B–C) Effect of the cytokinin signaling inhibitor S4893 co-treatment with CF during CIM culture on shoot regeneration. *: *p* < 0.05 in Dunnet’s test against mock treatment (*n* > 13). (D) Expression of SCR in CIM callus co-treated with S4893 and CF. (E) Expression of procambium-related genes in CIM callus under VLCFA-deficiency. *: FDR < 0.05 in edgeR. (F) Expression of fluorescence reporters for cambium genes in CF-treated CIM callus. (G–H) Shoot regeneration phenotypes of *lbd* mutants treated with CF during CIM culture. *: *p* < 0.05 in *t*-test (*n* > 27). (I) Schematic view of the role of VLCFA-mediated inter-layer communication in pluripotency acquisition. Representative images in (A, D, F) are from observation of at least 5 explants. Bars: 1 mm (B, G), 100 μm (A, D, F).

To test whether cytokinin hyperactivation is responsible for the defect in pluripotency acquisition, we performed co-treatment with CF and a cytokinin signaling inhibitor S4893 (*40*) on CIM. As expected, S4893 co-treatment suppressed ectopic TCSn:GFP expression caused by CF treatment (fig. S8B). We also verified that S4893 treatment did not affect CF-dependent decrease in VLCFA amount (fig. S2A). Strikingly, co-treatment with S4893 alleviated CF-induced defect in pluripotency acquisition. The co-treated callus after 5–6 days of CIM pre-incubation regenerated a comparative number of shoots with mock-treated callus (Fig. 6B, C and fig. S8C), with proper localization of SCR in the middle cell layers at CIM5D (Fig. 6D). These data clearly show that the misexpression of middle-cell layer-specific genes under CF treatment is caused by elevated cytokinin signaling.

We then tested whether VLCFAs regulate cytokinin synthesis or signaling. A previous report showed that cytokinin biosynthesis genes such as adenosine-phosphate isopentenyltransferase (*IPT*) *3/5/7* were transcriptionally upregulated upon VLCFA deprivation, leading to excessive cytokinin accumulation in seedlings (*14*). Our measurements reproduced accumulation of isopentenyladenine-type cytokinin in CF-treated seedlings (fig. S9A). In VLCFA-deficient callus, however, endogenous cytokinin levels were not elevated by CF treatment (fig. S9B). Consistent with the phytohormone quantification data, the overall expression of *IPT*s or other cytokinin biosynthesis genes was not upregulated (fig. S9C). These results indicate that the effect of VLCFA deprivation on cytokinin homeostasis is context-dependent, and it does not affect cytokinin biosynthesis in CIM callus. We therefore assumed that the hyperactivation of cytokinin signaling in VLCFA-deficient callus might instead result from an elevated response to exogenous cytokinin. To test this hypothesis, we performed CF treatment on cytokinin-free CIM. Under this condition, CF treatment no longer suppressed shoot regeneration (fig. S9D, E). Moreover, TCSn:GFP signal was not enhanced, and SCR localization remained normal even in the presence of CF (fig. S9F). We therefore conclude that the regeneration defect caused by VLCFA deficiency is attributable to an enhanced sensitivity to exogenous cytokinin.

Finally, we asked how elevated cytokinin signaling impairs pluripotency acquisition. Cytokinin signaling is generally known to inhibit lateral root development (*41*). More recently, cytokinin was shown to promote the fate transition of XPP cells toward cambium development rather than lateral root formation (*42*). During this process, cytokinin suppresses root stem cell regulators *WOX5* and *PLETHORA 1/2* (*PLT1/2*) through activation of cytokinin-responsive procambium and cambium regulators including LATERAL ORGAN BOUNDARIES DOMAIN 3/4 (LBD3/4) and WOX4 (*42*, *43*). As callus develops from XPP cells through a lateral-root-like genetic pathway (*3*, *4*), we hypothesized that a similar mechanism might underlie the defects caused by VLCFA deficiency. Consistent with this idea, genes associated with lignin and phenylpropanoid biosynthesis pathways were upregulated in the VLCFA-deficient calli in our RNA-seq analysis (Fig. 4A). Procambium regulator genes previously shown to be upregulated in cytokinin-treated lateral root primordia (*42*) were also induced in VLCFA-deficient calli (Fig. 6E). Our imaging analyses showed that the expression of a cork cambium marker *PEROXIDASE15* (*PER15*) expanded to the callus region under CF treatment (Fig. 6F). The reporter activity of *LBD4* also became stronger and spatially broader, occasionally extending into the middle-cell layer (Fig. 6F). Similar to TCSn:GFP, this CF-dependent expansion of the *LBD4* expression domain does not occur in the absence of exogenous cytokinin (fig. S9F). These observations suggest that VLCFA deprivation enhances the expression of procambium regulators in the callus through elevated cytokinin signaling.

To determine whether this expansion of procambium fate is responsible for the shoot regeneration defect, we treated mutants of cytokinin-inducible cambium *LBD* genes (*43*) with CF and analyzed their shoot regeneration phenotypes. *lbd4* single, *lbd4 lbd11* double, and *lbd1 lbd3 lbd4 lbd11* quadruple mutants exhibited reduced shoot regeneration (Fig. 6G, H), as previously reported for *lbd4* (*44*). Strikingly, we found that the *lbd1 lbd3 lbd4 lbd11* quadruple mutant was insensitive to CF treatment and produced a comparable number of shoots with the mock condition (Fig. 6G, H). This result clearly demonstrates that the regeneration defects caused by VLCFA deficiency depend on cytokinin-inducible procambium regulators. Based on these findings, we conclude that VLCFAs promote pluripotency acquisition by non-cell autonomously suppressing cytokinin signaling, which would otherwise promote procambium fate and inhibit establishment of the QC-like genetic network in the middle-cell layer.

## DISCUSSION

A key event in development of the multicellular body is the specification of cellular identity according to spatial patterns. Pluripotent callus is derived from a single cell layer and develops into a stratified structure composed of three distinct domains with specific gene expression profiles: the outermost, middle, and inner cell layers (*3*, *5*, *6*). It is well established that the middle-cell layer is essential for pluripotency acquisition. However, little was known about the patterning mechanisms of these domains. In this study, we show that VLCFA synthesis in the outermost callus layer plays essential roles for pluripotency acquisition and subsequent shoot regeneration. During pre-incubation on CIM, VLCFAs or their derivatives promote callus maturation by non-cell autonomously confining cytokinin response and the expression of procambium genes to inner layers. VLCFAs thereby allow the establishment of middle-cell layers with QC-specific gene expression. Based on these observations, we propose a model in which inhibitory relationships among these layer-specific regulators establish distinct spatial domains in the CIM callus (Fig. 6I). Our study highlights the intricate balancing mechanism among hormonal signals that underlie cellular pluripotency. Whereas exogenous auxin is the primary trigger for pluripotent callus formation, cytokinin supplemented in CIM promotes pluripotency by preventing fate determination toward root development, as evidenced by enhanced root production in calli precultured on CIM with S4893 or without CK (fig. S8C, S9D). On the other hand, excessive cytokinin signaling promotes a procambium-like fate and suppresses root identity gene expression (Fig. 6). It is therefore important for the outermost cells to synthesize VLCFAs, which spatially restrict cytokinin responses and thereby enable the establishment of the middle-cell layer.

VLCFAs reportedly have pleiotropic regulatory roles in root development and CIM callus formation. For instance, VLCFAs were reported as negative regulators of callus development (*16*, *17*), which may appear to contradict our findings that VLCFAs promote callus maturation. Those previous studies focused on the increased callus initiation foci along the longitudinal axis of uncut seedling roots upon VLCFA deficiency (fig. S10), which has been linked to polar auxin transport, rather than to cytokinin (*16*, *17*, *45*). By contrast, the present study focuses on gene expression patterning along the radial axis (fig. S10). Together, these studies reveal two separable modes of VLCFA-mediated control in callus development, namely, initiation and radial patterning. Our current study on callus radial patterning provides key insights into the role of VLCFAs on lateral root development (*16*, *18*, *19*) as well. For example, the VLCFA-deficient *puchi* mutant displays delayed lateral root development, yet the specific cause of the growth retardation remains to be identified. Considering that cytokinin signaling is up-regulated in *puchi* lateral root primordia (*46*), it is plausible that the elevated cytokinin signaling might also be the cause of lateral root phenotype of *puchi* mutant (*16*). Importantly, our genetic analyses argue against the possibility that VLCFAs regulate callus tissue patterning via production of cuticle (Fig. 2), which is essential for lateral root emergence (*47*). Our functional dissection of downstream pathways in this study is an important step toward elucidating the developmental roles of VLCFAs.

A recent study (*23*) implicated ATML1 together with VLCFAs in callus pluripotency, although it was not formally tested whether ATML1 is functionally involved in pluripotency acquisition. Our results instead showed that regulatory roles of VLCFAs in pluripotency acquisition is at least partly independent of ATML1/PDF2. Our phenotypic analyses with the *atml1 pdf2* double mutant and the inducible chimera repressor line ATML1-SRDX showed that ATML1/PDF2 are dispensable for pluripotent callus formation, while they play important roles in shoot morphogenesis on SIM. A previous study using *Nicotiana tabacum* discussed mutual regulatory relationships between NtML1 and NtWUS in establishing the spatial patterning (*48*). However, we did not detect any ectopic WUS expression in the outermost layer of shoot progenitors in *atml1 pdf2* mutant (Fig. 5C). We therefore conclude that ATML1/PDF2 play pivotal roles at later stages of shoot morphogenesis.

Our findings have broad implications beyond plant development. We demonstrate that a transient epidermis-like tissue on pluripotent callus plays a pivotal role in *de novo* plant organ regeneration, paralleling the critical role of a “wound epidermis” during organ regeneration in amphibians and planarians (*49–51*). Our findings therefore provide insights into the shared principles across kingdoms, whereby coordinated interactions between tissue layers guide the reconstruction of highly organized multicellular structures. VLCFAs and VLCFA-containing sphingolipids play important roles in cell fate determination in animals as well (*52*, *53*). In mammals, VLCFAs are involved in the epidermal stem cell differentiation (*53*). Future studies are awaited to test whether analogous mechanisms underlie VLCFA-mediated regulatory processes in plants and animals.

## MATERIALS AND METHODS

### Plant materials and culture conditions

*Arabidopsis thaliana* L. (Arabidopsis) accession Columbia (Col-0) was used as WT unless otherwise specified. Wassilewskija (Ws) was used along with Col-0 for comparison with the *atml1-1* and *pdf2-1* mutants, which were originally created in Ws background and backcrossed to Col-0 for at least three times (*35*). Landsberg *erecta* (Ler) was used for comparison with *cer4-1* (CS34)(*32*). All the other mutants listed below have the Col-0 background: *kcs1-5* (SALK_200839C)(*17*), *cer10-2* (SALK_088645)(*27*), *cer2-5* (SALK_084443)(*29*), *wax2* (SALK_020265)(*31*), *pas2-1* (CS73026)(*28*), *pas1-4* (SALK_051324)(*17*), *lbd4c*, *lbd4c lbd11c*, and *lbd1c lbd3c lbd4c lbd11c* (*43*). We used the following published transgenic lines: gWUS-3xGFP (*54*), pWOX5:ER-GFP (*55*), pSCR:GFP-SCR *scr-3* (*56*), gATML1-GFP (*22*), pATML1:NLS:3xGFP (*57*), proRPS5A:ATML1-SRDX/pER8 in pATML1:NLS:3xGFP (*37*), DR5rev:GFP (*58*), pPIN1:PIN1-YFP (*59*), TCSn:GFP (*38*), pPER15:erYFP (*60*, *61*), and pLBD4:erYFP (*43*). We introgressed gWUS-3xGFP, pWOX5:ER-GFP, pSCR:GFP-SCR, and TCSn:GFP to *kcs1-5* by crossing. Similarly, gWUS-3xGFP and pWOX5:ER-GFP were introgressed to *atml1-1 pdf2-1* by crossing.

Plants were grown on half-strength Murashige-Skoog (MS) medium containing 0.6% (w/v) Gelzan and 1% (w/v) sucrose at 22°C. For tissue culture experiments, 8–9-mm hypocotyl explants from 7DAS etiolated seedlings, 1 cm root explants from 7DAS seedlings grown under continuous light, or 7DAS seedlings grown under continuous light were placed on CIM [Gamborg B5 medium containing 0.25% (w/v) Gelzan, 2% (w/v) glucose, 2,4-D (0.5 mg/L), and kinetin (0.1 mg/liter)] under constant light at 22°C. For phenotyping of *pas2-1* (Fig. 3d–e), shorter (c.a. 6 mm) explants were used for both wildtype and the mutant because of impaired hypocotyl growth in this mutant. Only root explants were used for phenotyping *pas1-4* (Fig. 3f–g) because this mutant exhibited even severer defect in etiolated hypocotyl growth. CIM without kinetin [Gamborg B5 medium containing 0.25% (w/v) Gelzan, 2% (w/v) glucose, and 2,4-D (0.5 mg/L)] was used for the analyses in Fig. S9. For shoot and root regeneration, the explants cultured on CIM for 4 d (unless otherwise stated) were further cultured on SIM [Gamborg B5 medium containing 0.25% (w/v) Gelzan, 2%(w/v) glucose, indole-3-acetic acid (0.15 mg/L), and 2-iPA (0.5 mg/L)] and RIM [Gamborg B5 medium containing 0.25% (w/v) Gelzan, 2%(w/v) glucose, and Indole-3-butyric acid (0.5 mg/L)], respectively, under constant light at 22°C. The number of regenerated shoots was counted after 10 or 11 d of SIM culture and explant images were taken after 14 d of SIM culture, unless otherwise stated.

For chemical treatment, dexamethasone (Dex) (Fujifilm Wako), estradiol (Est) (Fujifilm Wako), cafenstrole (CF) (Fujifilm Wako), metazachlor (Meta) (Sigma-Aldrich), and S-4893 (EN300-761386; Enamine) were dissolved in ethanol (for Dex) or DMSO (for Est, CF, Meta, and S-4893) and supplied to the MS, CIM, or SIM media. Mock-treated controls received the same volume of the corresponding solvent.

### Imaging analyses

Images of explants were taken using a dissection microscope MZ10F equipped with a CCD camera DFC310FX (Leica). Confocal microscope SP8 (Leica) was used for cellular-level analyses. For confocal microscopy, samples were fixed in 4% (w/v) paraformaldehyde solution buffered by phosphate-buffered saline (PBS). The fixed samples were washed in PBS and then treated in ClearSee (*62*) solution containing 1% [v/v] Calcofluor White Stain solution (Sigma-Aldrich) for at least 1 d to clear the tissue and stain the cell wall. The samples were mounted on glass slides with ClearSee solution and observed with 405 nm (Calcofluor) and 488 nm (GFP and YFP) laser excitation. Images were processed and analyzed using Fiji (*63*). For tissue sectioning, samples were fixed in formalin acetic acid-alcohol fixative (5% [v/v] formalin, 5% [v/v] acetic acid, and 63% [v/v] ethanol), followed by treatment in a graded series of ethanol from 70% to 100%. The samples were then embedded in Technovit 7100 resin (Kulzer, Wehrheim, Germany), sectioned into 6 μm-thick sections by using a rotary microtome (RM 2165; Leica), and stained by toluidine blue. The sections were imaged using a light microscope (BX53M; Evident).

### Plant transformation and analyses of transgenic plants

We PCR-amplified 3.2 kb *KCS1* promoter (*17*) and *KCS1* coding sequence (CDS) fragments by using the following primers : KCS1pro_F1, GGGGACAACTTTGTATAGAAAAGTTGGAGTTAGCCAATAGCCATATGCG; KCS1pro_R2, GGGGACTGCTTTTTTGTACAAACTTGTCAGTATAGTTTTGGGTCGAAATATTTC; KCS1cds_F1, GGGGACAAGTTTGTACAAAAAAGCAGGCTATGGAGAGAACAAACAGC, KCS1cds_R, GGGGACCACTTTGTACAAGAAAGCTGGGTCTTGCACAACTTTAACCGG. The promoter and CDS fragments were cloned into pDONRp4-p1R and pDONR-221 vectors, respectively, by Gateway BP cloning system (Invitrogen). The promoter and CDS were then transferred to the binary vector R4pGWB504 (*64*) via an LR reaction (Invitrogen). The resultant binary vector was introduced into *Agrobacterium tumefaciens* strain GV3101 by electroporation and introduced to Arabidopsis *kcs1-5* plants following the floral dip method (*65*). T1 seeds were sown on selection media containing hygromycin and cefotaxime and cultured under the dark condition. After 7-day culture, transformed seedlings with elongated hypocotyls were used for the two-step tissue culture. We followed the same protocol as described above, except that we used CIM and SIM supplemented with cefotaxime to prevent bacterial growth. Some explants were fixed at CIM4D to observe KCS1-GFP signal. Regeneration phenotype was assessed at SIM21D.

### Cytokinin quantification

Approximately 70 mg of seedlings and hypocotyl explants were collected into a sampling tube at 4 d after transfer to CIM, frozen by liquid nitrogen, and homogenized by Tissue Lyser II (QIAGEN). Extraction and quantification of cytokinins were performed according to the method described previously (*66*, *67*). Endogenous cytokinins were measured by UPLC–ESI–qMS/MS using a UPLC system coupled to a tandem quadrupole mass spectrometer (ACQUITY Premier UPLC and Xevo TQ-XS; Waters). Chromatographic separation was performed on a pentafluorophenyl column (ACQUITY Premier HSS PFP, 1.8 μm, 2.1 × 150 mm, VanGuard FIT; Waters). Data were processed using MassLynx with TargetLynx 4.2 (Waters).

### VLCFA quantification

Approximately 30 mg of hypocotyl explants were collected into a sampling tube at 4 d after transfer to CIM, frozen by liquid nitrogen, and homogenized by Tissue Lyser II (QIAGEN). Pentadecanoic acid (C15:0) was added as an internal standard, and total lipids were extracted using the Bligh and Dyer method. Fatty acid methyl esters (FAMEs) were prepared by incubating the lipid extracts in methanolic HCl (Tokyo Chemical Industry Co., Ltd.) at 85°C for 90 min and were then extracted with hexane. FAMEs were quantified by GC-FID using an Agilent 8860 GC System equipped with an ULBON HR-SS-10 column (30 m, 0.25 mm i.d.; Shinwa Chemical Industries, Ltd.). Nitrogen was used as the carrier gas at a constant linear velocity of 20 cm s⁻¹. The injector temperature was set to 250°C. The oven temperature was initially set to 160°C, increased to 210°C at 2.1°C min⁻¹, and then held at 210°C for 1 min. The detector temperature was set to 250°C.

### RNA-seq

Hypocotyl explants were harvested for RNA extraction at 4 d after transfer to CIM. Total RNA was extracted using the RNeasy Plant Mini Kit (QIAGEN) according to the manufacturer’s protocol, with on-column DNase digestion (QIAGEN) to eliminate genomic DNA contamination. mRNA was isolated from 1 μg of total RNA using a Dynabeads mRNA DIRECT Micro Kit (Invitrogen) and used for cDNA library preparation with the NEBNext Ultra Directional RNA Library Prep Kit II for Illumina (New England Biolabs) following the manufacturer’s protocols. Briefly, polyadenylated mRNA was isolated directly from total RNA using Dynabeads Oligo(dT)25 (Invitrogen). mRNA was fragmented in the NEBNext First Strand Synthesis Reaction Buffer by heating at 94°C for 15 min. First-strand cDNA was synthesized from the fragmented mRNA by reverse transcription and subsequently used as a template for second-strand cDNA synthesis. During second-strand synthesis, deoxythymidine triphosphate was replaced with deoxyuridine triphosphate (dUTP). The cDNA was end repaired, dA-tailed, and ligated to the NEBNext Adaptor. The second-strand cDNA containing dUTP was digested with USER enzyme, and sequencing tags and barcodes were introduced by 12 cycles of PCR amplification. The amplified libraries were purified using AMPure XP beads (Beckman Coulter). Library quality was assessed using the TapeStation system with the High Sensitivity D1000 ScreenTape assay (Agilent Technologies). RNA sequencing was performed using 150-bp paired-end reads on an Illumina HiSeq X Ten platform. The reads were trimmed using Fastp (v1.0.1)(*68*), after which they were aligned to the Arabidopsis transcriptome (TAIR10) using RSEM (v1.3.1) with Bowtie2 (*69*). Differentially expressed transcripts were identified using edgeR package (v3.40.2) in R/Bioconductor with with FDR < 0.05 and |logFC| > 0.5 cutoffs. GO analysis was performed using the clusterProfiler package (4.6.2)(*70*) of R (v4.2.2).

### Statistical analyses

Statistical tests were performed using R (v4.2.2) with significance threshold of *p* < 0.05 unless otherwise specified.

## Acknowledgments

We thank Tatsuaki Goh (NAIST), Keiji Nakajima (NAIST) and Shinobu Takada (Osaka University) for providing plant materials; Naoko Hisanaga (NAIST) and Yuriko Ikeda (NAIST) for technical assistance; Hidehiro Fukaki (Kobe University) and Akihito Mamiya (Kyoto University) for constructive comments. RNA-seq experiments and confocal microscopy observation were supported by the NAIST Life Science Collaboration Center (LiSCo).

## Author contributions

YD and MI conceived the project and designed the experiments. YD performed most of the experiments and analyses. TN quantified fatty acids. MK and HS quantified phytohormones. KMI and TM prepared libraries for RNA-seq. KN, MA, and APM generated some of the plant materials and provided critical comments. YD and MI wrote the manuscript with input from all authors.

## Funding

This work was supported by JSPS KAKENHI (JP23KJ1579 and JP24H00710 to YD; JP24K09489 to TN; JP26K21757 to HS and MK; 25K02297, 25H02564 and 26H01727 to MI), the JST FOREST Program (JPMJFR214H to M.I.), and the Research Council of Finland (grant number 364181 to APM).

## Competing interests

The authors declare no competing interests.

## Data availability

Sequence read data generated for the RNA-seq experiments will be available in the NCBI Sequence Read Archive (SRA).

---

**Supplementary Figure S1.**
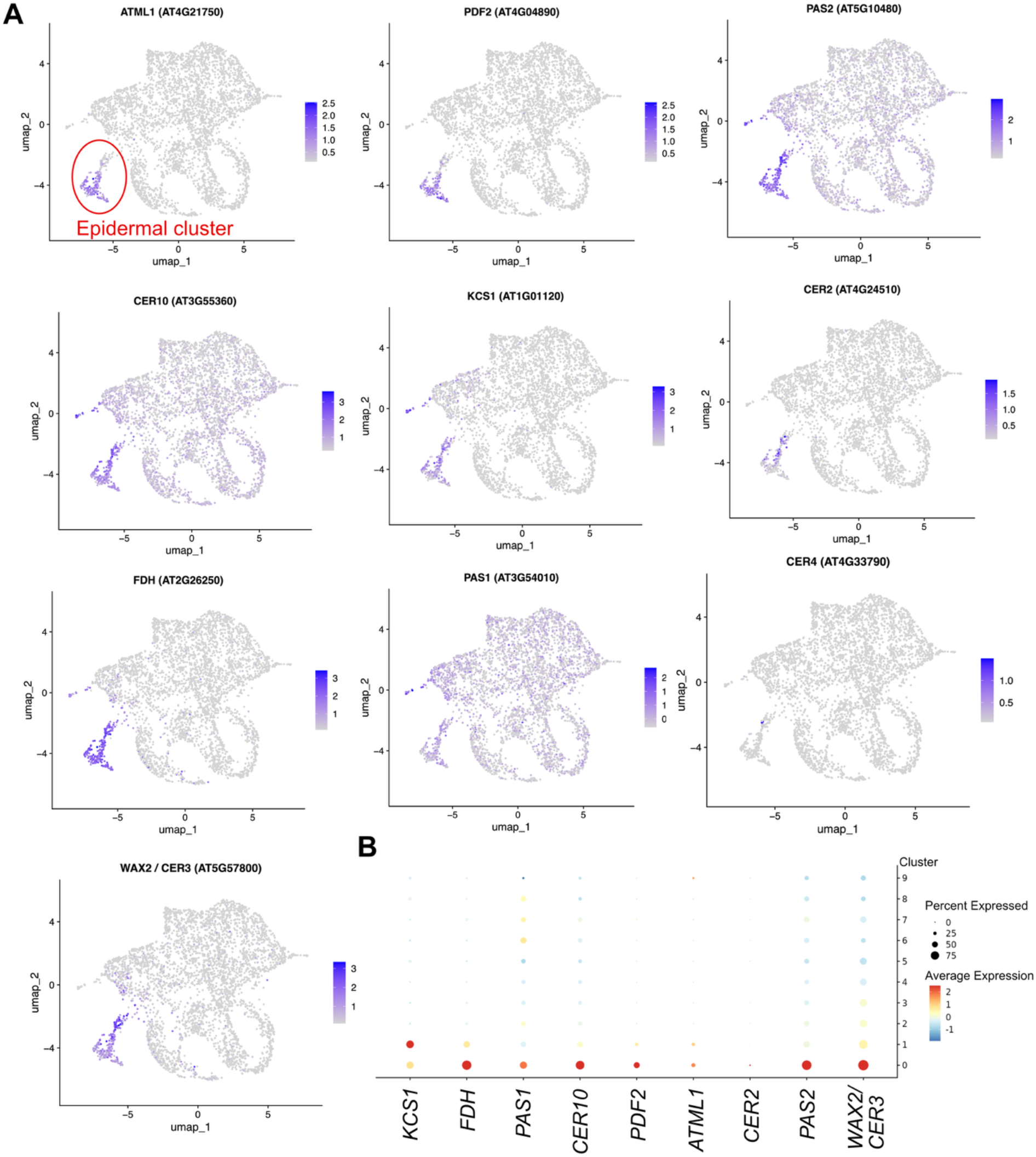
Expression of ATML1/PDF2, VLCFA-related genes, and cuticular wax synthesis genes in previous single cell RNA-seq datasets. (A) Data from SIM7D callus (*7*). Epidermal cell cluster (Cluster 9) marked by *ATML1/PDF2* expression is indicated by the red circle. (B) Data from CIM6D callus (*5*). Clusters 0 and 1 represent the outer layer of the callus marked by *ATML1/PDF2* expression.

**Supplementary Figure S2.**
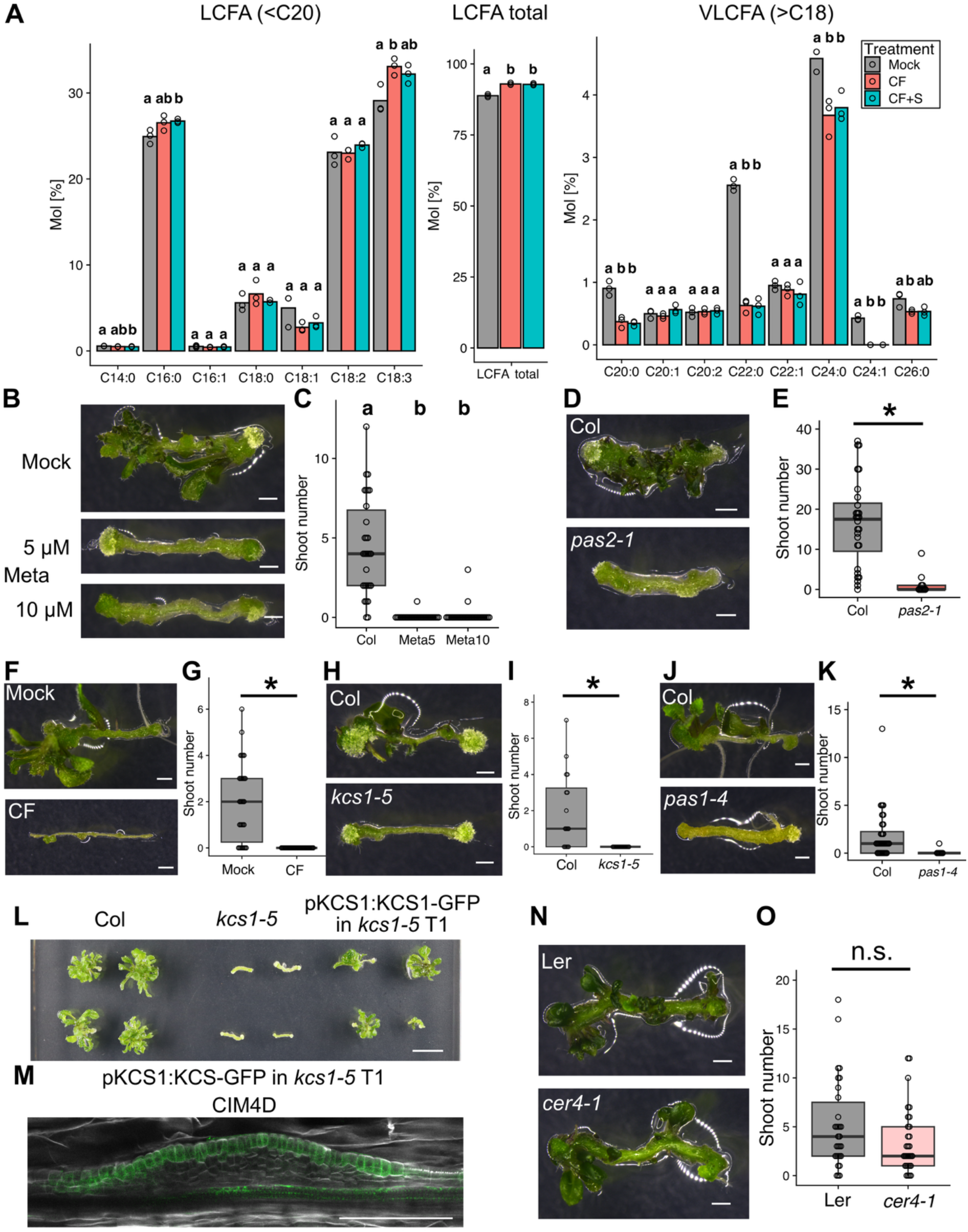
VLCFA synthesis is necessary for shoot regeneration from hypocotyl or root explants. (A) Total fatty acid composition of CF-treated and S4893- and CF-cotreated callus at CIM4D. Different letters indicate statistically significant differences (*p* < 0.05 in Tukey-Kramer test, *n* = 4). (B–C) Shoot regeneration phenotype of wild-type hypocotyl explants under continuous metazachlor (Meta) treatment. Different letters indicate statistically significant differences. (*p* < 0.05 in Tukey-Kramer test, *n* = 30). (D–E) Shoot regeneration phenotype of *pas2-1* hypocotyl explants. (*n* = 10 for *pas2-1* and *n* = 28 for Col). (F–G) Shoot regeneration phenotype of wild-type root explants under continuous CF treatment. (*n* = 30) (H–I) Shoot regeneration phenotype of *kcs1-5* root explants. (*n* = 16). (J–K) Shoot regeneration phenotype of *pas1-4* mutant root explants. (*n* = 7 for *pas1-4* and *n* = 32 for Col). (L–M) Complementation of the *kcs1-5* mutant phenotype in shoot regeneration from hypocotyl explants. (N–O) Shoot regeneration phenotype of *cer4-1* hypocotyl explants. The result of *t*-test is shown (*p* = 0.11, *n* = 30). M: Representative images from observation of at least 5 explants are shown. Bars: 1 mm (B, D, F, H, L, N), 100 μm (M). * : *p* < 0.05 in *t*-test.

**Supplementary Figure S3.**
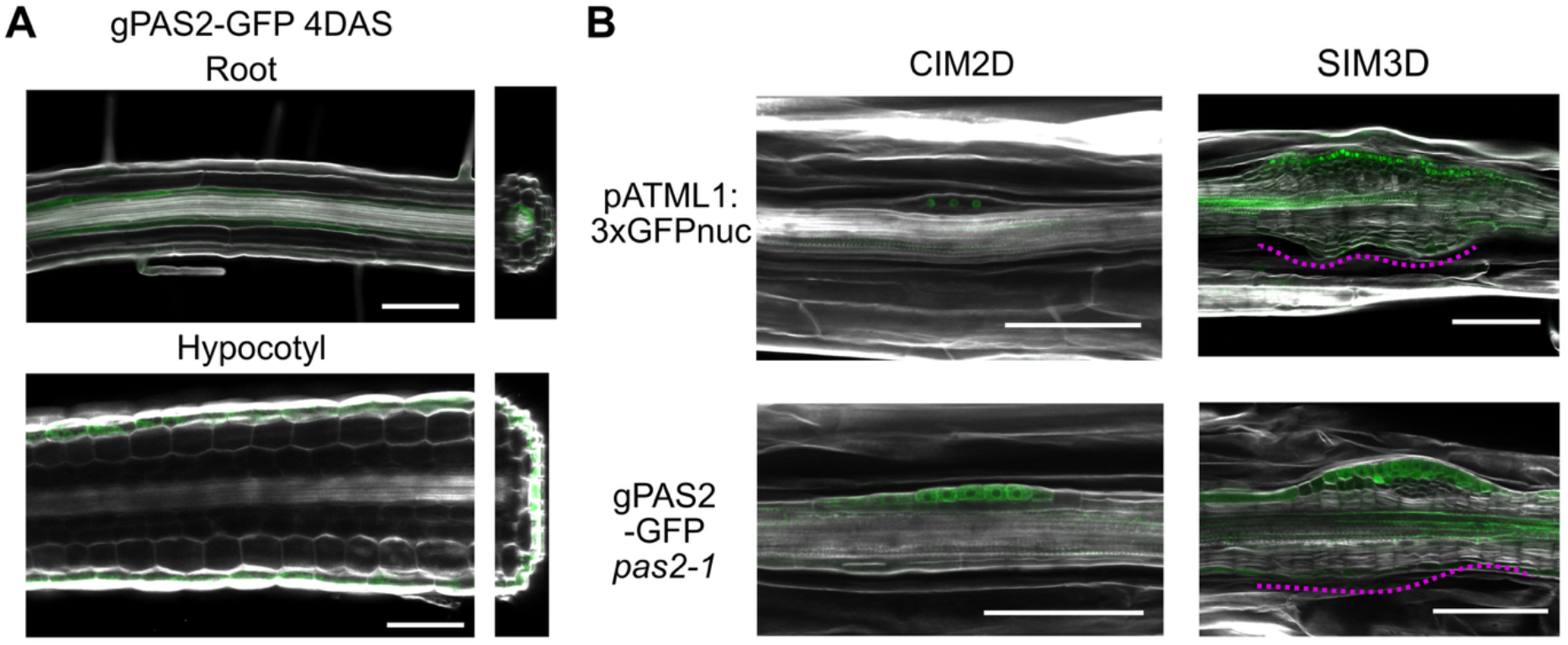
Localization of fluorescence reporters for *ATML1* and PAS2. (A) Localization of PAS2-GFP in the light-grown seedling root and hypocotyl. Reconstructed cross-section images are shown in the right. (B) Localization of reporters during two-step culture. Dotted magenta lines indicate callus regions without epidermal marker expression. Baras: 100 μm.

**Supplementary Figure S4.**
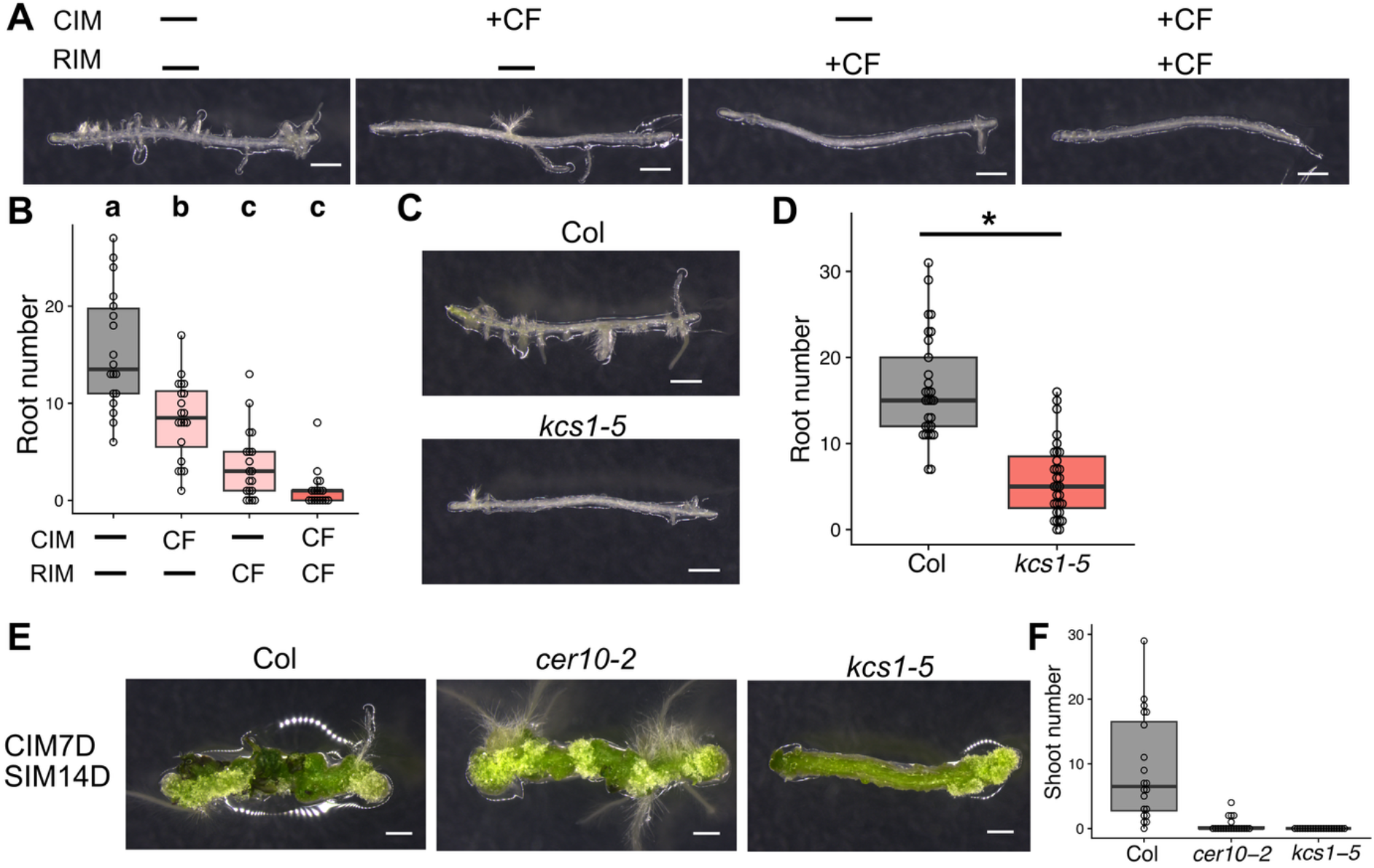
VLCFA synthesis is necessary for pluripotency acquisition of CIM callus. (A–B) Effect of stage-specific CF treatment on adventitious root regeneration from wild-type hypocotyl explants. Different letters indicate statistically significant differences (*p* < 0.05 in Tukey-Kramer test, *n* > 16). (C–D) Adventitious root regeneration phenotype of *kcs1-5* hypocotyl explants. *: *p* < 0.05 in *t*-test (*n* > 28). (E–F) Effect of prolonged CIM culture on shoot regeneration of VLCFE-deficient mutant callus. Shoot number was counted on SIM14D. *n* = 20. Bars: 1 mm.

**Supplementary Figure S5.**
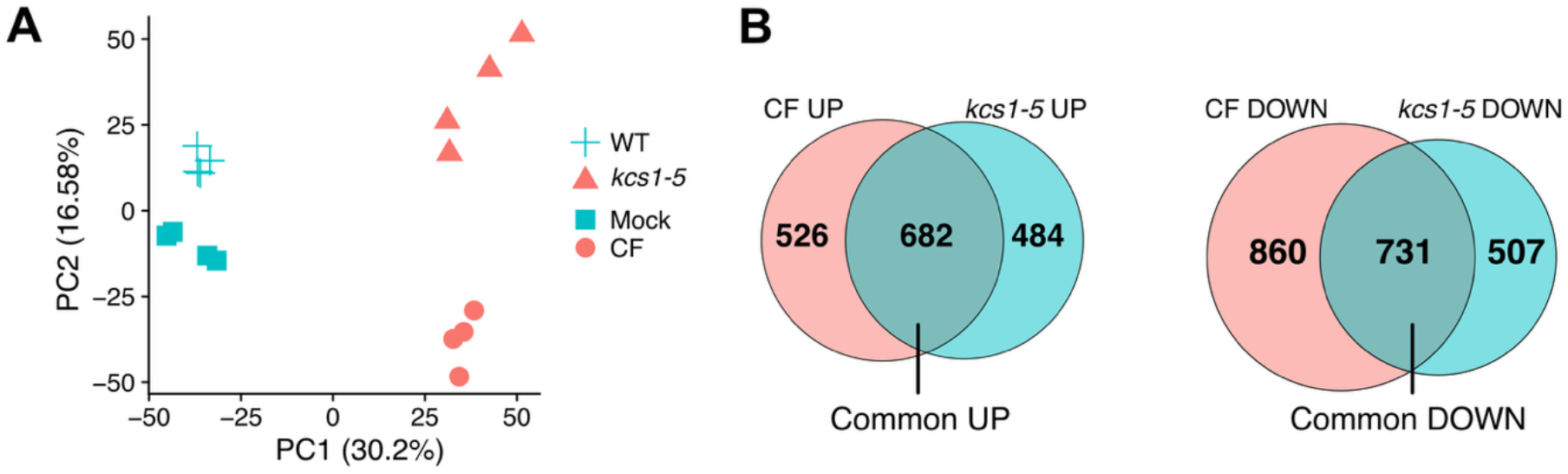
RNA-seq analysis of VLCFA-deficient callus. (A) PCA plot of analyzed samples (B) Venn diagram showing the number of DEGs for the two VLCFA-deficient callus samples.

**Supplementary Figure S6.**
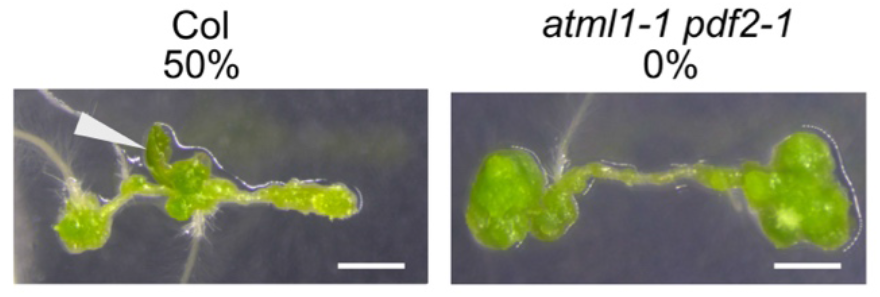
Function of ATML1/PDF in regeneration from root explants. Shoot regeneration phenotype of *atml1-1 pdf2-1* double mutant root explants. The frequency of shoot regeneration is shown (*n* = 14 (Col), 11 (*atml1-1 pdf2-1*)). Arrowheads indicate regenerated shoots. Bars: 1 mm.

**Supplementary Figure S7.**
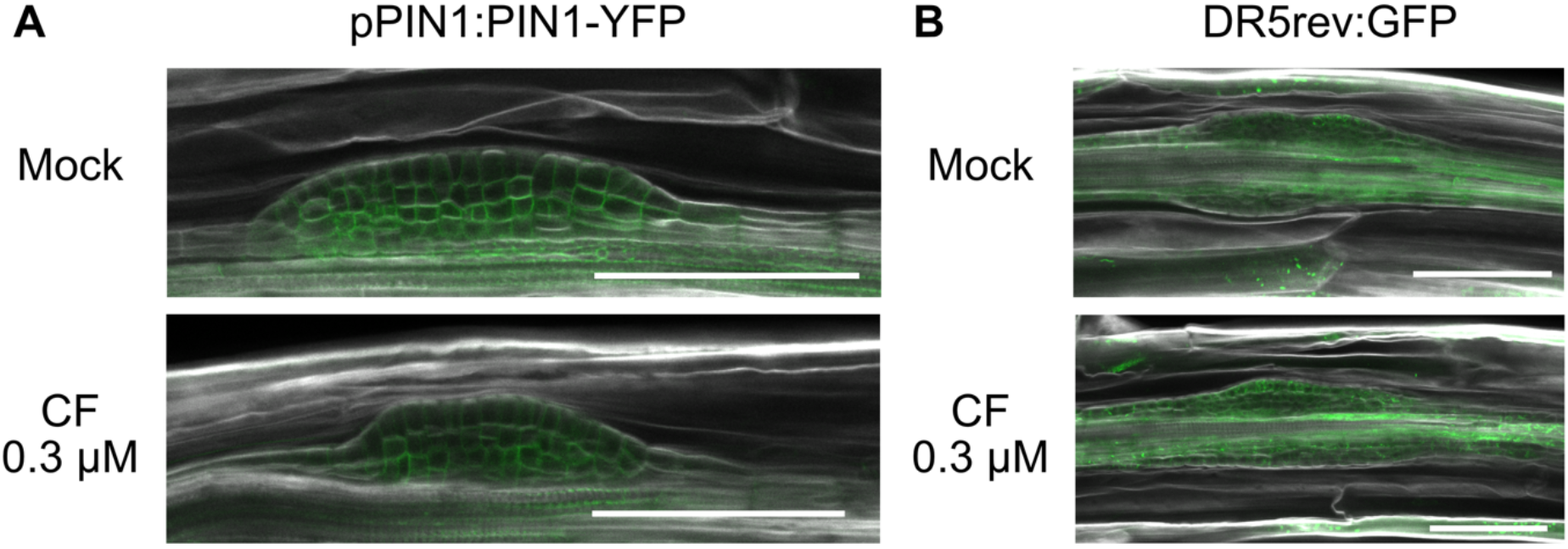
VLCFA deprivation in CIM callus may not substantially alter auxin dynamics. (A) Localization of pPIN1:PIN1-YFP in CIM callus treated with CF. (B) Expression of auxin response reporter DR5rev:GFP in CIM callus treated with CF. Representative images from observation of at least 5 explants are shown. Bars: 100 μm.

**Supplementary Figure S8.**
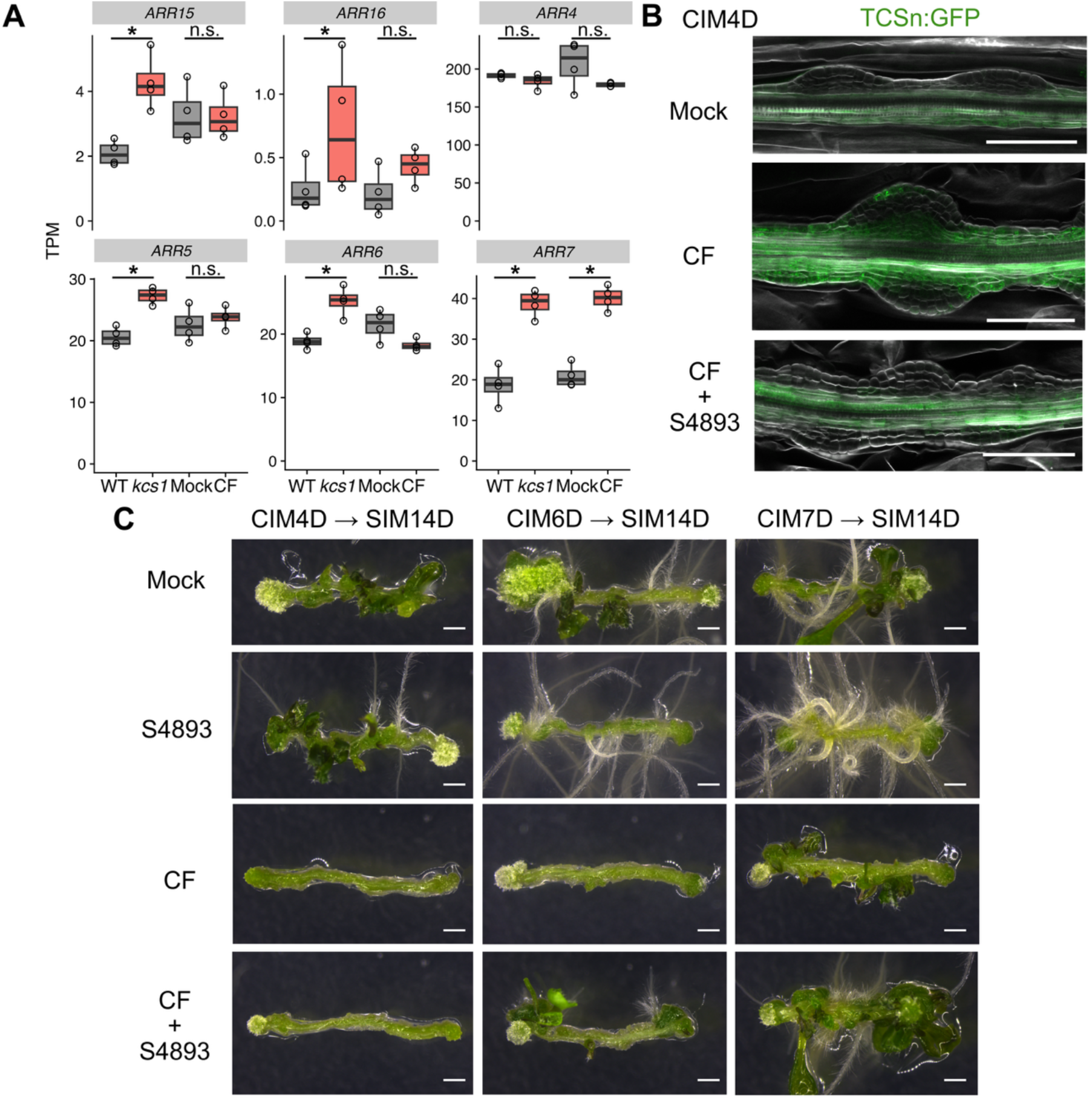
VLCFAs regulate pluripotency acquisition by suppressing cytokinin signaling. (A) Expression of cytokinin-responsive type-A *ARR* genes in VLCFA-deficient CIM callus. *: FDR < 0.05 in edgeR. (B) Expression of cytokinin response marker TCSn:GFP in CIM callus under cotreatment with CF and S4893. Representative images from observation of at least 5 explants are shown. (C) Effect of the cytokinin signaling inhibitor S4893 co-treatment with CF during CIM culture on shoot regeneration. See Fig. 6C for quantification results. Bars: 1 mm (C), 100 μm (B).

**Supplementary Figure S9.**
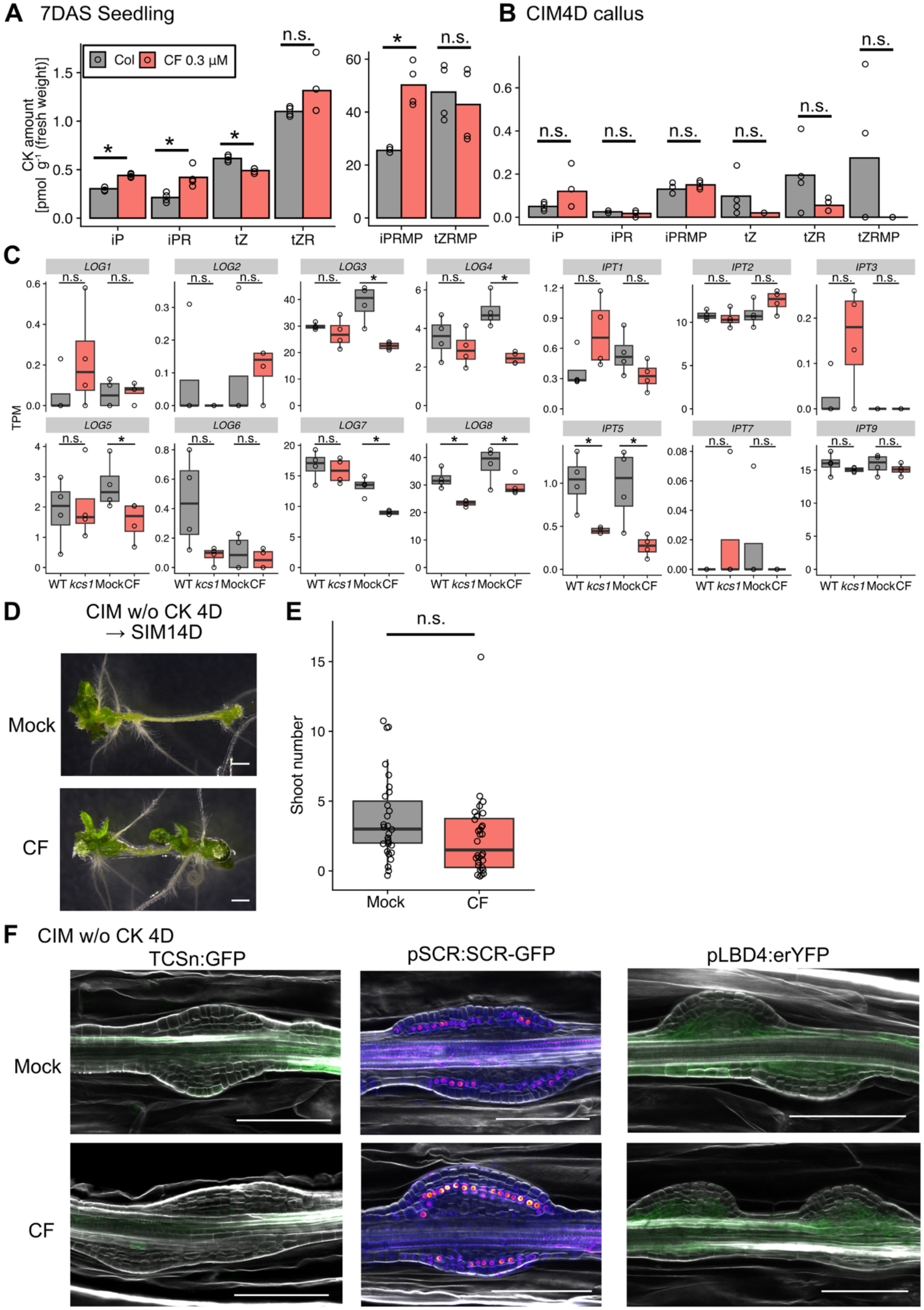
VLCFAs regulate cellular responsiveness to exogenous cytokinin. (A, C) Endogenous cytokinin content in 7 DAS seedling and callus at CIM4D under CF treatment. iP, *N^6^*-(Δ^2^-isopentenyl)adenine; iPR, iP riboside; iPRMP, iPR 5’-monophosphate; tZ, *trans*-zeatin; tZR, tZ riboside; tZRMP, tZR 5’-monophosphate. (B) Expression of cytokinin biosynthesis genes in VLCFA-deficient CIM callus. *: FDR < 0.05 in edgeR. (D–F) Effect of CF treatment during preculture on cytokinin-free CIM. (D–E) Shoot regeneration phenotype under CF treatment. The result of *t*-test is shown (*p* = 0.12, *n* > 28). (F) Expression of cytokinin response marker (TCSn:GFP), pluripotency marker (SCR-GFP), and cambium-related *LBD4* reporter in CF-treated callus. Representative images from observation of at least 5 explants are shown. Bars: 1 mm (C), 100 μm (E).

**Supplementary Figure S10.**
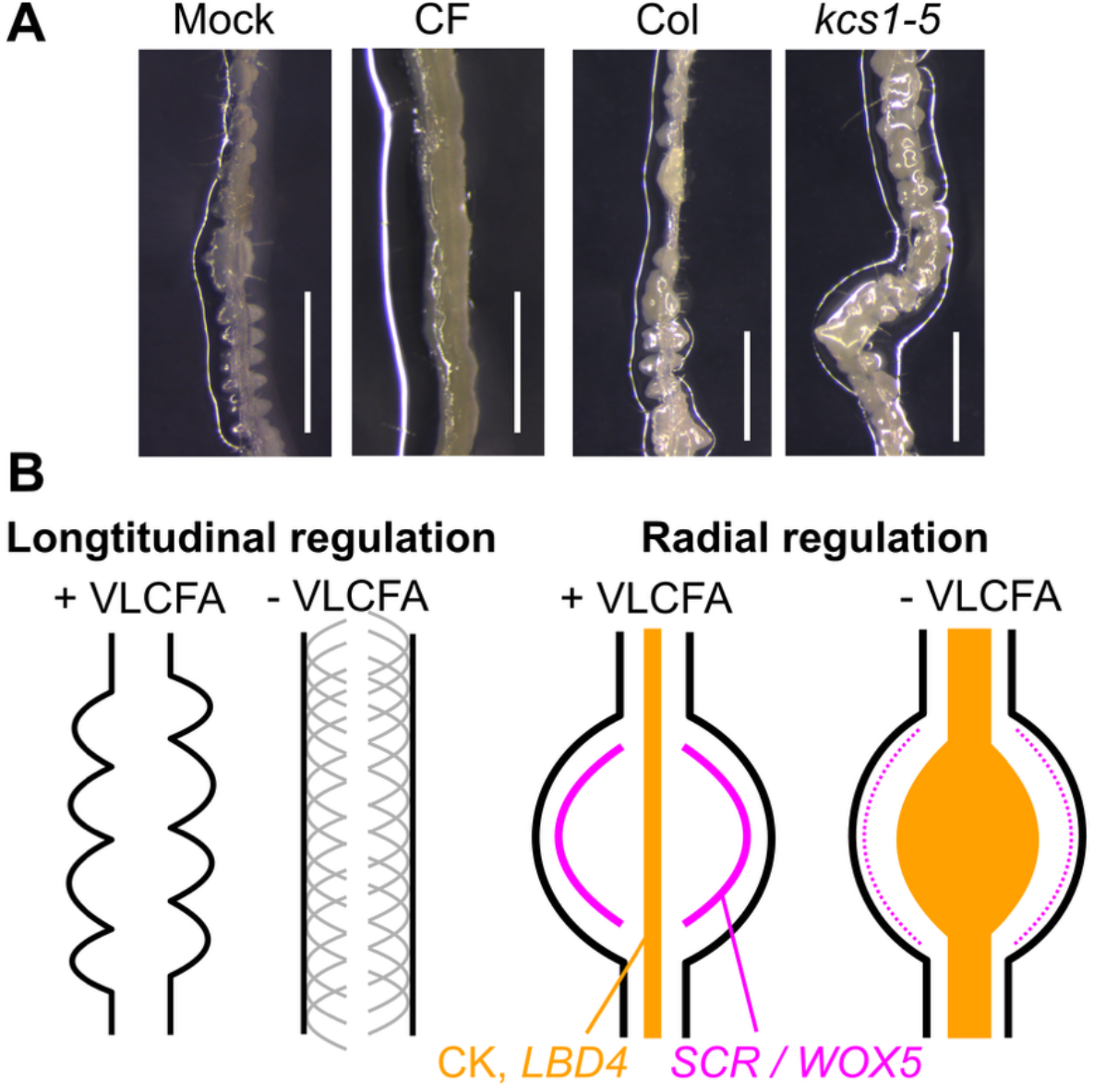
Effect of VLCFA-deficiency on callus growth of seedling roots. (A) Phenotypes of seedling roots placed on CIM for 5 days. Bars: 1 mm. (B) Schematic views of distinct VLCFA functions.

## Notes

### Competing Interest Statement

The authors have declared no competing interest.

